# A novel exopolysaccharide pathway from a freshwater *Sphingomonas* isolate

**DOI:** 10.1101/2023.11.03.565537

**Authors:** Alexandra G. Goetsch, Daniel Ufearo, Griffin Keiser, Christian Heiss, Parastoo Azadi, David M. Hershey

## Abstract

Bacteria embellish their cell envelopes with a variety of specialized polysaccharides. Biosynthesis pathways for these glycans are complex, and final products vary greatly in their chemical structures, physical properties and biological activities. This tremendous diversity comes from the ability to arrange complex pools of monosaccharide building blocks into polymers with many possible linkage configurations. Due to the complex chemistry of bacterial glycans, very few biosynthetic pathways have been defined in detail. To better understand the breadth of polysaccharide production in nature we isolated a bacterium from Lake Michigan called *Sphingomonas sp.* LM7 that is proficient in exopolysaccharide (EPS) production. We identified genes that contribute to EPS biosynthesis in LM7 by screening a transposon mutant library for colonies displaying altered colony morphology. A gene cluster was identified that appears to encode a complete *wzy/wzx-*dependent polysaccharide assembly pathway. Deleting individual genes in this cluster caused a non-mucoid phenotype and a corresponding loss of EPS secretion, confirming that LM7 assembles a novel *wzy/wzx*-dependent polysaccharide. We extracted EPS from LM7 cultures and showed that it contains a linear chain of 3- and 4-linked glucose, galactose, and glucuronic acid residues. Finally, we found that the EPS pathway we identified diverges from those of adhesive polysaccharides such as the holdfast that are conserved in higher *Alphaproteobacteria.* Our approach of characterizing complete biosynthetic pathways holds promise for engineering of polysaccharides with valuable properties.

**Importance:** Bacteria produce complex polysaccharides that serve a range of biological functions. These polymers often have properties that make them attractive for industrial applications, but they remain woefully underutilized. In this work, we studied a novel polysaccharide called promonan that is produced by *Sphingomonas sp.* LM7, a bacterium we isolated from Lake Michigan. We extracted promonan from LM7 cultures and identified which sugars are present in the polymer. We also identified the genes responsible for polysaccharide production. Comparing the promonan genes to those of other bacteria showed that promonan is distinct from previously characterized polysaccharides. We conclude by discussing how the promonan pathway could be used to produce new polysaccharides through genetic engineering.

## Introduction

Bacterial cells are decorated with elaborate arrays of carbohydrates (1). Polysaccharides such as capsules, O-antigens and exopolysaccharides (EPSs) serve as a physical interface between the surface of the bacterium and other objects in the environment (2, 3). These glycans have a tremendous influence on fitness, and their properties have specialized to reflect the wide range of niches colonized by bacteria (4). Immunomodulators (5), adhesins (6) and gelling agents (7) are among the many classes of bacterial polysaccharides that have been identified. Collectively, these polymers present a rich source of chemical diversity that can be utilized for human benefit.

Bacteria synthesize complex polysaccharides using the *wzy* strategy. This process initiates with glycosyltransferase (GT) enzymes that attach monosaccharides sequentially onto a lipid carrier at the cytoplasmic face of the inner membrane, producing a glycolipid called the repeating unit. A highly conserved set of assembly proteins then flips (Wzx) repeating units across the cytoplasmic membrane, polymerizes (Wzy) them into a chain and secretes (Wzc, Wza) the mature glycan into the extracellular space (8). The sugar composition and linkage configurations present in *wzy*-dependent polysaccharides vary widely. Each biosynthetic pathway utilizes a unique set of GTs to assemble a repeating unit oligosaccharide containing 3-8 sugars (9). Variability in substrate and linkage specificities among GT enzymes gives rise to the vast chemical diversity of bacterial glycans.

Bacterial polysaccharides have the potential to be used in a range of applications, but few have found widespread use. Current production pipelines rely on extracting polysaccharides directly from producing organisms (10–12). This approach restricts the potential of glycans that are produced at lower quantities or by organisms that are not amenable to large scale fermentation. We are interested in developing an alternative approach whereby bacteria are genetically modified to produce polysaccharides recombinantly. Not only would this strategy allow for over-production of low abundance glycans and facilitate the characterization of biosynthesis pathways from genetically intractable organisms, but it could also provide a framework for engineering novel polysaccharides through combinatorial biosynthesis. For instance, the sugar compositions of natural glycans could be modified by exchanging genes for GTs with different substrate specificities. Such engineering efforts require a pool of glycan assembly enzymes with well-characterized specificities that can be used to introduce chemical complexity.

Biosynthetic pathways for *wzy-*dependent polysaccharides have seldom been characterized in detail. Capsular polysaccharide A from *Bacteroides fragilis* (13), succinoglycan from *Sinorhizobim meliloti* (14), xanthan gum from *Xanthomonas campestris* (15) and colonic acid from *Escherichia coli* (16) represent rare examples for which the enzymatic steps of repeating unit assembly have been clearly defined. For each of these glycans, a clear biomedical, industrial or agricultural relevance provided justification for the considerable effort required to elucidate the biosynthetic pathways. However, these model pathways encompass only a miniscule fraction of the chemical diversity seen in bacterial glycans (17), and polysaccharides with promising properties continue to be identified (18–20). Defining additional pathways for *wzy-*dependent polysaccharides will lead to the discovery of novel enzymatic functionalities.

To explore the breath of polysaccharide biosynthesis in nature, we began isolating bacteria from freshwater lakes that were proficient in polysaccharide production. We describe here an isolate from Lake Michigan called *Sphingomonas sp.* LM7. LM7 displayed a mucoid colony morphology on agar plates that was indicative of EPS production. We utilized a genetic screen based on colony morphology to identify genes involved in polysaccharide biosynthesis. Two genetic loci emerged as key determinants of colony morphology. One locus contains genes for producing a secreted polysaccharide that we have named promonan, and the other contains genes for producing a second polysaccharide that remains associated with the cell surface. We defined the monosaccharide composition of the promonan EPS, performed a detailed analysis of genes involved in its biosynthesis and compared its biosynthetic pathway to those of polysaccharides produced by related bacteria. Our results indicate that the promonan EPS belongs to a novel family of *wzy-*dependent polysaccharides that could serve as a valuable scaffold for producing engineered glycans.

## Results

### *Isolation and phenotypic characterization of Sphingomonas sp.* LM7

We collected water from Lake Michigan at Promontory Point in Chicago, IL and used it to isolate bacteria by serial dilution on PYE agar (21). Isolate LM7 was selected for further study based on its colony morphology on agar plates. The organism grew as “mucoid” yellow colonies that display a shiny, gelatinous appearance (Fig 1A). This mucoid phenotype was more pronounced when the bacterium was incubated at lower temperatures and on agar plates supplemented with sugars such as glucose or sucrose. We predicted that LM7 was secreting a novel polysaccharide because the mucoid colony morphology is often associated with exopolysaccharide (EPS) production.

**Figure 1:**
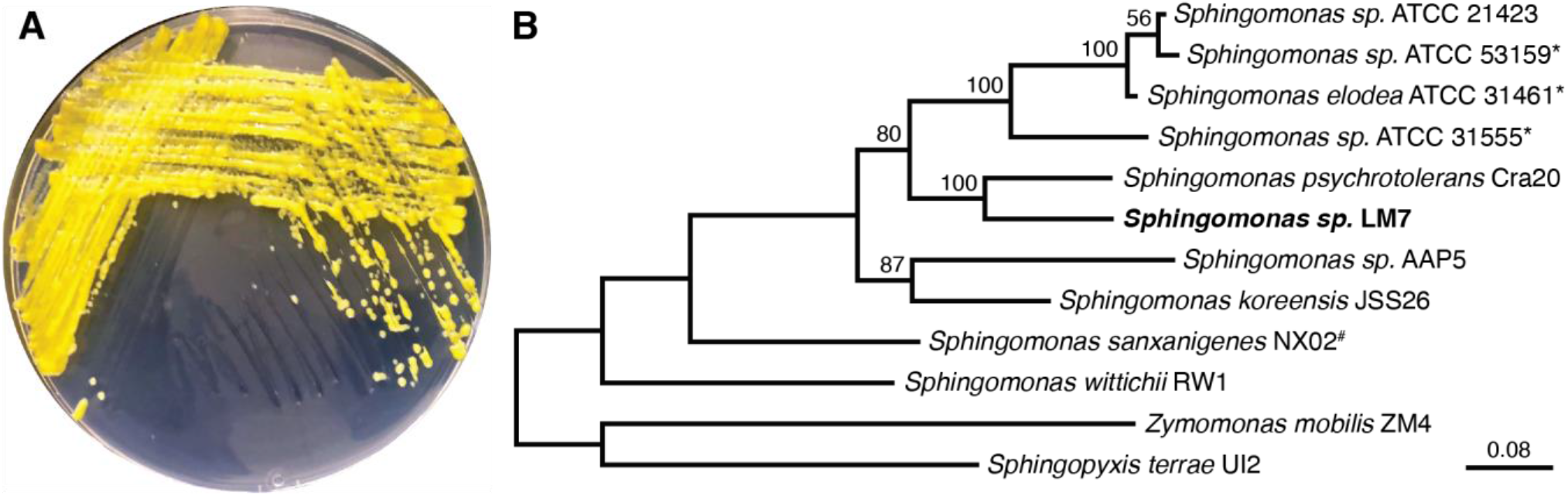
Isolation of *Sphingomonas sp.* LM7. (A) *Sphingomonas sp.* LM7 growing on agar plates. The isolate grows as yellow, mucoid colonies. (B) Phylogenetic tree showing select isolates from the *Sphingomonadales* order of *Alphaproteobacteria.* The neighbor-joining tree was constructed from 16S rDNA sequences aligned with ClustalX. Bootstrapping values are indicated at the relevant nodes. * indicates canonical sphingan producing strains. ^#^ highlights the non-canonical sphingan called sanxan produced by *Sphingomonas sanxanigenes* NX02. Note that the sanxan biosynthesis genes appear distinct from the sphingan or promonan genes (56).

Sequencing of the 16S rRNA gene placed LM7 in the genus *Sphingomonas,* with the most similar pure culture isolate being *Sphinogmonas psychrotolerans* Cra20 (95% identity) (22). Certain members of the genus *Sphiongomonas* produce EPSs known as sphingans (gellan, welan, diutan) (23), and we were curious if LM7 was producing a sphingan-type polysaccharide. We sequenced the genome of the LM7 and found that it contains a single, circular chromosome of 4,155,514 bases. There are two 16S rRNA genes with identical sequences. We constructed a phylogenetic tree based on the 16S rRNA gene sequences of several isolates within the order *Sphingomonadales.* LM7 and *S. psychrotolerans* form a clade that is distinct from canonical sphingan producing strains (Figure 1B). We were also unable to identify loci in the LM7 genome that resemble the *gel* cluster of genes for sphingan biosynthesis (24). We predicted that *Sphingomonas sp.* LM7 was producing a novel EPS that is distinct from the previously characterized sphingan family.

### Identification of genes affecting colony morphology

We performed a genetic screen to study the molecular basis for EPS production in LM7. A transposon mutant library was plated on a defined medium supplemented with glucose. The overwhelming majority of transposon insertion mutants show the mucoid phenotype under these conditions. We screened the mutants for colonies that have a non-mucoid, “matte” appearance (Fig 2A). 250 individual mutants with altered colony morphology were isolated, and the transposon insertion sites were mapped by arbitrary PCR. The insertion sites are summarized in Table 1.

**Figure 2:**
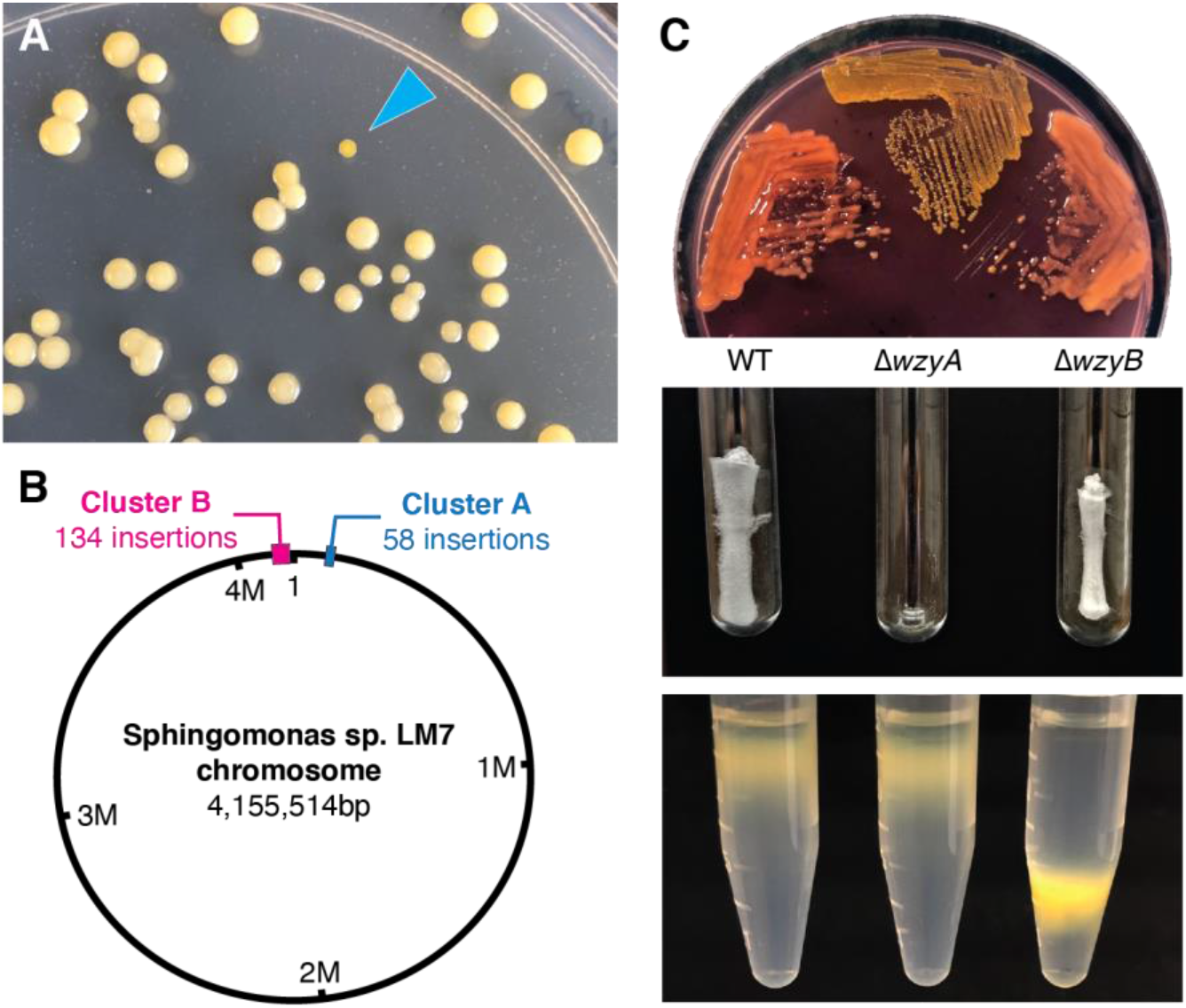
Identification of two polysaccharide biosynthesis clusters in LM7. (A) Screening for transposon insertion mutants with altered colony morphology. Mutagenized cells were plated on agar plates. Under these conditions, the majority of colonies show the mucoid morphology. The blue carat points to a non-mucoid mutant. (B) Map of the LM7 genome showing the locations of the two gene clusters that influence colony morphology. (C) Phenotypes of Δ*wzyA* and Δ*wzyB* mutants. The top image shows growth of LM7 strains on solid medium containing Congo Red dye. Wild type and the Δ*wzyB* mutant appear mucoid and react with the Congo Red dye, while the Δ*wzyA* mutant does not. The middle image shows an EPS extraction. Wild-type and Δ*wzyB* cultures produce secreted matrix. Δ*wzyA* cells do not secrete this matrix. The bottom image shows density gradient centrifugation of LM7 strains. The Δ*wzyB* mutant displays increased cell density relative to wild type and Δ*wzyA* cells.

**Table 1:** Sites of transposon insertions affecting colony morphology in LM7.

| Gene | Frequency | Annotation |
| --- | --- | --- |
| RS00295 | 1 | PAS domain-containing methyl-accepting chemotaxis protein |
| RS00350 | 7 | WcaJ family polyisoprenylphosphate hexose-1-phosphate transferase (PHPT) |
| upstream RS00360 | 1 | Predicted promoter region of BXU08_RS00360 |
| RS00360 | 4 | Beta-barrel outer membrane porin |
| RS00365 | 2 | Wza family outer-membrane polysaccharide export (OPX) protein |
| RS00370 | 2 | AAA+ family ATPase |
| RS00375 | 4 | Wzz family polysaccharide chain length determinant |
| RS00380 | 3 | P-loop NTPase |
| RS00385 | 4 | WecG/TagA family glycosyltransferase |
| RS00395 | 5 | Wzy family polysaccharide polymerase |
| RS00400 | 1 | GT2 family glycosyltransferase |
| RS00405 | 1 | GT2 family glycosyltransferase |
| RS00410 | 6 | GT2 family glycosyltransferase |
| RS00415 | 2 | GH10 family glycosylhydrolase |
| RS00420 | 9 | OafA family acyltransferase |
| RS00425 | 4 | GT2 family glycosyltransferase |
| RS00430 | 10 | GH16 family glycosylhydrolase |
| RS00665 | 2 | DedA family membrane protein |
| RS02105 | 2 | PAS domain-containing sensor histidine kinase |
| RS06730 | 1 | Tryptophan halogenase |
| RS07760 | 1 | Type II secretion system protein GspL |
| RS07770 | 1 | Type II secretion system protein GspH |
| upstreamRS09000 | 1 | Predicted promoter region of BXU08_RS09000 |
| RS09000 | 2 | OmpR family response regulator |
| RS10090 | 1 | Hypothetical protein |
| RS10945 | 1 | Acetoacetyl-CoA reductase PhbB |
| RS11025 | 2 | Poly(R)-hydroxyalkanoic acid synthase PhaC |
| RS12915 | 1 | Helix turn helix family protein |
| RS13595 | 1 | FliA family transcriptional regulator |
| RS14850 | 1 | bifunctional (p)ppGpp synthase-hydrolase SpoT |
| RS15820 | 1 | UDP-Galactose 4-epimerase GalE |
| RS18205 | 1 | GT2 family glycosyltransferase |
| RS18930 | 2 | Major facilitator superfamily (MFS) transporter |
| upstreamRS19155 | 1 | Predicted promoter region of BXU08_RS19155 |
| RS19155 | 3 | Bifunctional GGDEF-EAL diguanylate cyclase-hydrolase |
| RS19160 | 6 | TPR domain containing sensor histidine kinase |
| downstreamRS19160 | 3 | Insertion downstream of BXU08_RS19160 |
| RS19205 | 1 | NAD(P)+-dependent nucleotide sugar dehydrogenase |
| RS19215 | 12 | Amidotransferase, exosortase associated |
| RS19220 | 18 | PLP-dependent aminotransferase |
| RS19225 | 8 | WxcM family acetyltransferase |
| RS19230 | 8 | MviM family dehydrogenase |
| RS19235 | 14 | GT4 family glycosyltransferase |
| RS19250 | 8 | SDR family NAD(P)<sup>+</sup>-dependent oxidoreductase |
| RS19260 | 12 | WbuX family N-acetyl sugar amidotransferase |
| RS19265 | 10 | AlgZ/HisF2 family acetamidino modification system |
| RS19270 | 9 | HisH family amidotransferase subunit |
| RS19290 | 2 | EpsG family glycosyltransferase |
| RS19295 | 12 | GT4 family glycosyltransferase |
| RS19300 | 2 | Wzy family polysaccharide polymerase |
| RS19310 | 2 | Wzc family tyrosine autokinase |
| RS19360 | 3 | NAD(P)<sup>+</sup>-dependent epimerase |

6 of the 250 mutants contained insertions in intergenic regions, while the remaining 244 mapped to coding regions. 48 open reading frames and 4 intergenic regions were disrupted. Insertion sites were scattered across the chromosome, but the majority of the mutations (192 of 250) were concentrated at two loci (Fig 2B). The first locus (Cluster A in Fig2B) contained 58 insertions. It spans ∼18.5kb and encodes a gene cluster that contains 17 open reading frames. The second locus (Cluster B in Fig2B) is separated from Cluster A by ∼102kb. 134 mutations were identified in Cluster B, which spans a nearly 48kb region encompassing over 40 open reading frames. We conclude that the genetic determinants of colony morphology in *Sphingomonas sp.* LM7 are concentrated at two separate loci in the genome.

### Two polysaccharide biosynthesis gene clusters

Each of the two loci that affect colony morphology in LM7 contains genes with homology to *wzy, wzx, wzz* and *wza* along with a suite of GT genes, suggesting that two distinct polysaccharides influence colony morphology under our growth conditions. To differentiate the roles of Cluster A and Cluster B in polysaccharide production we generated in-frame deletions (Δ*wzyA* and Δ*wzyB*) of the genes predicted to encode the Wzy polymerase enzyme from each cluster. We found that including Congo Red dye in agar plates accentuated the mucoid phenotype by causing wild-type LM7 colonies to take on a reddish tint. The Δ*wzyA* mutant displayed a rough texture on Congo Red plates and remained yellow in appearance, indicating that this mutant did not produce a Congo Red reactive matrix. The Δ*wzyB* mutant appeared indistinguishable from the wild-type strain on Congo Red plates. The colonies remained mucoid and took on the reddish hue of the Congo Red dye (Fig 2C). These results indicate that LM7 produces a Congo Red reactive matrix that requires the *wzyA* gene but not the *wzyB* gene.

LM7 secretes a gelatinous matrix into liquid medium during growth that can be precipitated with ethanol. This material is resistant to treatment with protease and DNAse and remains in the aqueous phase during a phenol fractionation. We subjected spent medium from the wild-type, Δ*wzyA* and Δ*wzyB* strains to an extraction that had been optimized for isolation of this secreted matrix. Lyophilized extracts prepared from wild-type LM7 cultures contained a white, fibrous material reminiscent of cotton. The fibrous material was completely absent in extracts prepared from the Δ*wzyA* mutant, but the Δ*wzyB* mutant produced extracts that were indistinguishable from wild-type. Thus, *wzyA,* but not *wzyB,* is required for secretion of the gelatinous material (Fig 2C).

Some of the transposon mutants isolated in our initial screen appeared to sediment differently during centrifugation. We predicted that differences in the appearance of cell pellets reflected changes in the density of the cell envelope. To investigate cell densities, we subjected our strains to density gradient centrifugation in a solution of Percoll. Wild-type cells remain near the top of the gradient under our centrifugation conditions, and the Δ*wzyA* mutant retained the buoyancy of the wild-type strain. Δ*wzyB* cells sediment further along the gradient than wild-type cells, indicating that this mutation causes the cell envelope to become denser (Fig 2C). We conclude that *wzyB* is required for producing a polysaccharide that increases the buoyancy of the LM7 cell envelope.

Our analysis of the Δ*wzyA* and Δ*wzyB* mutants is consistent with a model in which LM7 produces two distinct polysaccharides. Most of the colony morphology mutants we isolated had transposon insertions in one of two regions of the genome, and each of these loci appears to encode a complete set of genes for producing a *wzy-*dependent polysaccharide. Deleting the *wzy* gene, which codes for the repeating unit polymerase enzyme, from the two gene clusters caused disparate phenotypes. The Δ*wzyA* mutation disrupted the LM7’s ability to secrete a gelatinous, Congo Red reactive EPS, while the Δ*wzyB* mutant displayed a denser cell envelope. We conclude that Cluster A is responsible for the production of a secreted polysaccharide and that Cluster B contains the genes for a cell surface-associated polysaccharide.

### Isolation and analysis of a secreted polysaccharide

We developed a method to purify the secreted EPS from LM7. Growth at lower temperatures (18°C) and supplementing the growth medium with glucose (2%) led to enhanced EPS secretion. EPS could be precipitated from spent medium of LM7 cultures with alcohol and subsequently enriched with a liquid-liquid extraction using aqueous phenol. After dialysis and lyophilization, the resulting extracts contained a patch of fluffy, white fibers. When the extraction was carried out with Δ*wzy* cultures, only trace amounts of an oily residue were observed (Fig 2C). Thus, our extraction leads to the purification of a secreted polysaccharide that requires the *wzyA* gene. We have named this polysaccharide promonan after the site on Promontory Point in Chicago, IL from which LM7 was isolated.

Promonan extracts were subjected to a series of chemical analyses to characterize the polysaccharide’s chemical composition. Analysis of *O-*trimethylsilyl (TMS) methyl glycoside derivatives prepared from promonan showed that glucose, glucuronic acid and galactose were the major monosaccharide constituents of promonan (Table 2). The exact ratios of the three sugars could not be determined from this analysis, but glucose was clearly more abundant than the other two residues. We studied the linkage patterns in promonan using a recently developed method for analyzing partially-methylated alditol acetate derivatives (PMAAs) derived from uronic acid sugars (25). All three monosaccharides (Glc, GlcA and Gal) were present in 3- and 4-linked forms (Table 3). Trace amounts (≤ 1% of total) of terminal and multiply linked forms of each monosaccharide were also detected. Our analysis indicates that promonan is composed of glucose, glucuronic acid and galactose. It also suggests that the repeating unit comprises a linear chain containing 3- and 4-linked forms of each residue.

**Table 2:** Monosaccharide composition of promonan EPS. Monosaccharides were identified by GC/MS analysis of trimethylsilane derivatives.

| Glycosyl Residue | Estimated mole percent |
| --- | --- |
| Glucuronic Acid (GlcA) | 23.9 |
| Galactose (Gal) | 16.5 |
| Glucose (Glc) | 59.6 |

**Table 3:** Linkage analysis of promonan EPS. Semi-quantitative analysis of partially-methylated alditol acetates (PMAAs) prepared from promonan extracts.

| Residue | Relative peak area (%) |
| --- | --- |
| Terminal Glucopyranosyl (t-Glc) | 1.0 |
| Terminal Glucopyranosyl Uronic Acid (t-GlcA) | 1.0 |
| Terminal Galactopyranosyl (t-Gal) | 0.9 |
| 3-linked Glucopyranosyl (3-Glc) | 10.6 |
| 3-linked Glucopyranosyl Uronic Acid (3-GlcA) | 10.9 |
| 3-linked Galactopyranosyl (3-Gal) | 4.0 |
| 4-linked Galactopyranosyl (4-Gal) | 8.8 |
| 4-linked Glucopyranosyl (4-Glc) | 32.1 |
| 4-linked Glucopyranosyl Uronic Acid (4-GlcA) | 30.4 |
| 3,4-linked Galactopyranosyl (3,4-Gal) | 0.1 |
| 3,4-linked Glucopyranosyl (3,4-Glc) | 0.2 |
| 3,6- linked Glucopyranosyl (3,6-Glc) | 0.1 |

### Targeted deletions of genes in the promonan cluster

The promonan locus (Cluster A) contains 17 predicted open reading frames (ORFs, BXU08_RS00350-BXU08_RS00430). 16 of the ORFs are arranged in the (+) direction on the chromosome, and a single ORF (BXU08_RS00430) sits at the 3’ end of the locus in the (-) direction (Fig 3A). ORFs predicted to encode a polyisoprenyl-phosphate hexose-phosphate transferase (PHPT), five GTs and a full suite of polysaccharide assembly/secretion factors (Wza, CapB, Wzz, Wzx, Wzy) make up a core set of polysaccharide biosynthesis genes. Additional genes for an outer membrane porin (OMP), a AAA+ family ATPase, an acyltransferase and two glycosylhydrolases are also present along with a small (64AA) ORF with no clear homology to any characterized protein families (Table 4). We predicted that this cluster contains the factors required for promonan biosynthesis and renamed these genes *prmA-Q* accordingly.

**Figure 3:**
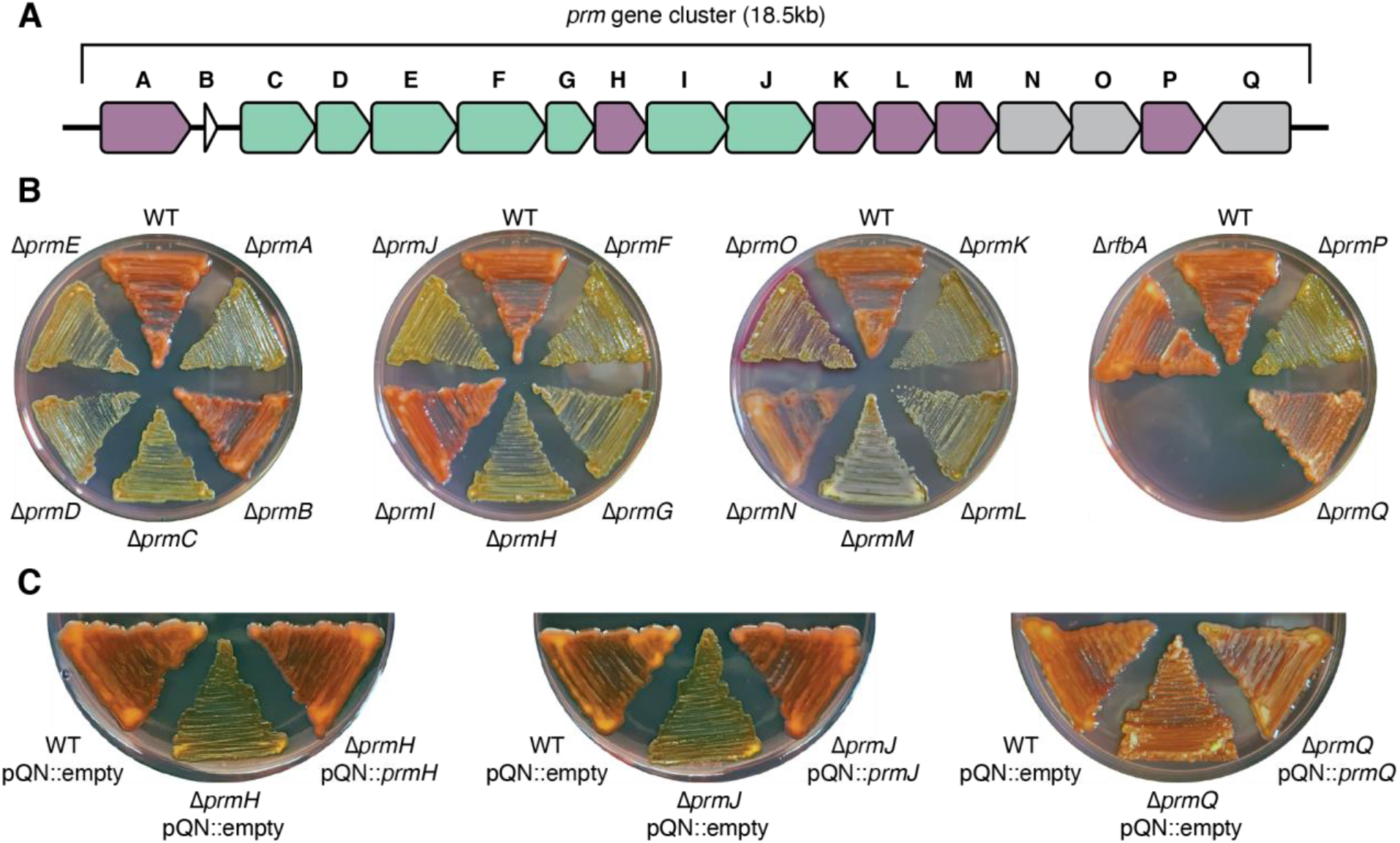
The promonan biosynthesis cluster. (A) Map of the *prm* cluster. Open-reading frames are colored by predicted function. Purple: monosaccharide incorporation; green: assembly/secretion; grey: polysaccharide modification. (B) Morphologies of individual *prm* mutants on solid medium supplemented with Congo Red. (C) Complementation of *prmH, prmJ* and *prmQ* deletion mutants. Non-mucoid phenotypes (Δ*prmH* and Δ*prmJ*) and the rugose phenotype (Δ*prmQ*) can be reverted by introducing the relevant gene *in trans*.

**Table 4:**
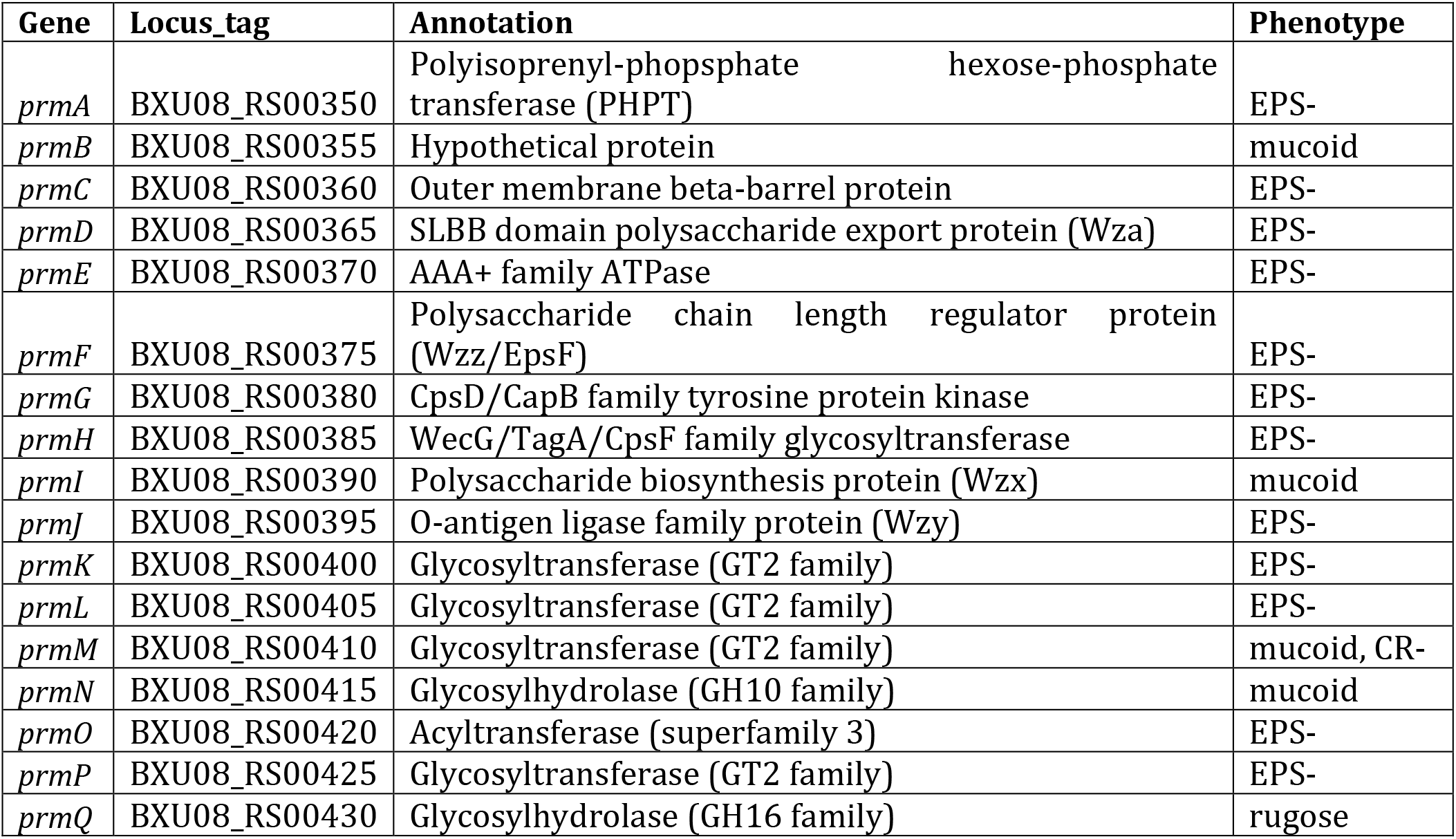
Functional annotations and mutant phenotypes for promonan biosynthesis genes.

We generated in-frame deletions for each of the 17 genes in the promonan cluster and assessed their EPS production phenotypes (Fig 3B). Nearly all of the deletions led to a loss of Congo Red staining and promonan secretion, suggesting that these mutations caused an EPS-phenotype. The Δ*prmB* (hypothetical), Δ*prmI* (*wzx*) and Δ*prmN* (GH10) mutations had no effect on colony morphology. Δ*prmM* (GT) cells displayed a mucoid phenotype on agar plates but did not react with the Congo Red dye. The Δ*prmQ* (GH16) mutant had a non-mucoid phenotype that we classified rugose because it could be distinguished from other non-mucoid mutants by its dry, crusty appearance and its ability to react with Congo Red.

The phenotypic effects of deleting genes in the promonan cluster were confirmed through ectopic complementation. We used a cumate inducible system developed by Kaczmarczyk, Vorholt and Francez-Charlot (26) to express *prm* genes from a plasmid in their respective deletion backgrounds. Figure 3C shows the complementation of the non-mucoid phenotypes in Δ*prmH* and Δ*prmJ* as well as the rugose phenotype in Δ*prmQ*. Taken together, our reverse genetic analysis confirms that the promonan cluster contains genes responsible for EPS production in LM7 and clarifies the role of each gene at this locus.

### Comparison of polysaccharide biosynthesis genes among Alphaproteobacteria

We compared the promonan EPS to polysaccharides produced by other *Alphaproteobacteria.* Certain isolates from the genus *Sphingomonas* produce acidic polysaccharides called sphingans that have gel-like properties reminiscent of our promonan extracts (27). The promonan gene cluster we identified does not show similarity to the conserved cluster for sphingan production (24). There is no recognizable synteny among core *wzy* genes (*wzy, wzx, wzy, wzz,* etc.). The promonan cluster lacks ABC transporter genes and genes for dTDP-rhamnose synthesis. Finally, the presence of a predicted WecG-type GT gene (*prmH*) and two glycosylhydrolases (*prmN* and *prmQ*), all three of which are not present in sphingan gene clusters, further distinguishes the promonan cluster from the sphingan genes.

The presence of rhamnose in the repeating unit is a hallmark of sphingan EPSs. We did not detect rhamnose in the purified promonan extracts and could not identify a complete set of genes for dTDP-rhamnose biosynthesis in the LM7 genome. Homologs of *rfbA* and *rfbB* are present. *rfbD* appears to be present, but it codes for a protein that is fused to an additional beta-galactosidase domain. We could not identify *rfbC* in the LM7 genome. Though our genomic analysis suggests that LM7 is likely incapable of producing dTDP-rhamnose, we confirmed that rhamnose is not incorporated into promonan by deleting *rfbA* (BXU08_RS19140). The resulting Δ*rfbA* mutant showed wild-type colony morphology and Congo Red staining (Fig 3B). Thus, the chemical compositions and genetic requirements for EPS production support the model that promonan and sphingan represent distinct EPS families.

Many *Alphaproteobacteria* assemble adhesive polysaccharides at their cell poles (28–32), of which the holdfast polysaccharide from *Caulobacter crescentus* has been studied most extensively (33). We were intrigued by the identification of a WecG-type glycosyltransferase gene (*prmH*) in the promonan cluster because a WecG homolog (HfsJ) is also required for holdfast production (34). WecG-family GTs are thought to catalyze the committed step in select polysaccharide biosynthesis pathways by attaching a second sugar onto a lipid-linked monosaccharide (35–37). We analyzed the genomes of a panel of *Alphaproteobacteria* that are known to produce holdfast-like adhesins. These bacteria, which span the phylogenetic diversity of *Alphaproteobacteria*, each contained a WecG-family GT gene (Fig 4A). We predicted that the WecG-family GTs we identified might participate in a conserved pathway for producing holdfast-like adhesins.

**Figure 4:**
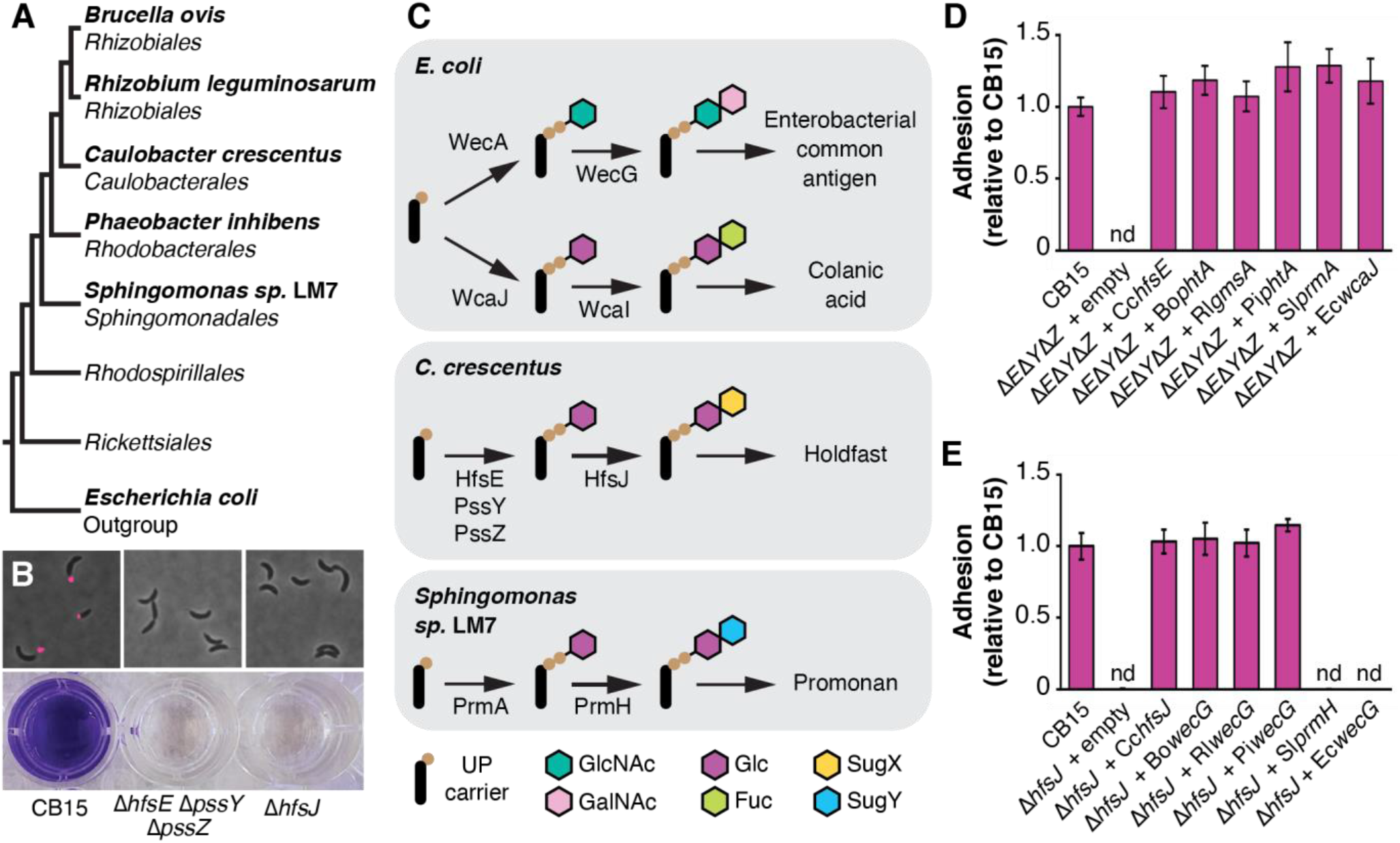
Conservation of polysaccharide biosynthesis genes in *Alphaproteobacteria*. (A) Schematic of *Alphaproteobacterial* phylogeny showing the relationship among isolates with putative polysaccharide adhesin pathways. (B) Phenotypes associated with holdfast production. fWGA stained shows the loss of holdfast production in the *C. crescentus* Δ*hfsE* Δ*pssY* Δ*pssZ* and Δ*hfsJ* mutants. Crystal violet (CV) staining of cultures grown in microtiter plates shows the loss of surface adhesion in the Δ*hfsE* Δ*pssY* Δ*pssZ* and Δ*hfsJ* mutants. (C) Inferred activities for initiation and subsequent glycosyltransferase enzymes in four polysaccharide pathways. (D) CV staining assay testing the effect of introducing PHPT genes from various bacteria into the *C. crescentus* Δ*hfsE* Δ*pssY* Δ*pssZ* mutant. All tested genes appear capable of converting UPP to UPP-Glc. (E) CV staining assay testing the effect of introducing *wecG*-family GT genes from various bacteria into the *C. crescentus* Δ*hfsJ* mutant. *wecGs* from *Brucella ovis, Rhizobium leguminosarum* and *Phaeobacter inhibens* can carry of the HfsJ reaction, while *wecG* from *Escherichia coli* (negative control) and *prmH* from *Sphingomonas sp.* LM7 cannot.

We tested the conservation of holdfast-like EPS pathways by performing a series of cross complementation experiments in *C. crescentus.* We reasoned that if an enzyme from one bacterium performs the same function as an enzyme in the holdfast pathway, introducing that gene into the corresponding *C. crescentus* deletion mutant should restore holdfast production. Polysaccharide biosynthesis begins when an initiator enzyme generates a lipid-linked monosaccharide. The production of undecaprenyldiphosphate-glucose (UPP-Glc) by the redundant action of HfsE, PssY and PssZ is thought to represent the initiation reaction in the holdfast pathway, and the product of this reaction likely serves as a substrate for HfsJ (38). Deleting the genes for these three initiators (Δ*hfsE* Δ*pssY* Δ*pssZ*) causes a holdfast null phenotype (39) that can be scored using a crystal violet-based adhesion assay and visualized by staining holdfasts with fluorescently labelled wheat germ agglutinin (fWGA, Fig 4B). The *wcaJ* gene from *Escherichia coli* codes for a undecaprenylphosphate glucose-phosphate transferase that produces UPP-Glc for colonic acid biosynthesis (38), and we confirmed that introducing *wcaJ* into the Δ*hfsE* Δ*pssY* Δ*pssZ* mutant restored holdfast production. PHPT genes from each of the *Alphaproteobacteria* in our panel, including *prmA* from LM7, also restored holdfast production to the Δ*hfsE* Δ*pssY* Δ*pssZ* mutant, indicating that each of these bacteria encodes an enzyme that can support holdfast synthesis by producing UPP-Glc (Fig 4D).

Deleting the gene for the WecG-family GT *hfsJ* causes a holdfast null phenotype in *C. crescentus* (34). HfsJ is thought to generate a lipid-linked disaccharide by adding an unknown sugar to UPP-Glc (produced by HfsE/PssY/PssZ). We introduced *wecG* homologs from our strain panel into the Δ*hfsJ* deletion to examine if they could restore holdfast production. *wecG*s from more derived (40) *Alphaproteobacterial* clades (*Rhizobiales, Caulobacterales, Roseobacterales*) complemented the holdfast defect in Δ*hfsJ,* but *prmH* from LM7 did not. The *wecG* gene from *E. coli* codes for an enzyme that utilizes a different lipid-linked acceptor substrate (UPP-GlcNAc) than *C. crescentus* HfsJ (UPP-Glc), and we confirmed that introducing *EcwecG* did not restore holdfast production to the Δ*hfsJ* mutant (Fig 4E). These results indicate that *wecG* homologs from representative *Rhizobiales, Caulobacterales* and *Roseobacterales* can convert UPP-Glc to a conserved UPP-linked disaccharide, but that PrmH from LM7 likely converts UPP-Glc to a different product. We conclude that a conserved pathway for holdfast-like adhesins exists in more derived clades of *Alphaproteobacteria* but that promonan production likely represents a distinct biosynthetic pathway.

## Discussion

The production of complex polysaccharides is nearly ubiquitous among bacteria. Despite the diversity of physical, chemical and biological properties encompassed by bacterial polysaccharides, they remain underutilized in industrial settings. Our long-term goal is to harness the vast chemical space occupied by these glycans for the development of polymers with valuable properties. To this end, we began isolating bacteria from environmental sources and dissecting their polysaccharide biosynthesis pathways. An isolate from Lake Michigan called *Sphingomonas sp.* LM7 secretes an exopolysaccharide (EPS) that we have named promonan. We identified a gene cluster that contains the factors required for promonan biosynthesis, defined the chemical composition of the polysaccharide and showed that promonan represents a novel family of EPSs.

Promonan biosynthesis clearly proceeds through a *wzy-*dependent mechanism. The *prm* locus contains genes for an initiating sugar phosphate transferase (*prmA*), five GTs (*prmH, prmK, prmL, prmM, prmP*) and a full set of polymerization/export factors (*prmD, prmF, prmG, prmI, prmJ*). *prmC* encodes an outer membrane porin (OMP) that likely also plays a role in polysaccharide export (41, 42). Thus, 12 of the 17 genes in the *prm* cluster make up a core set of factors that should be sufficient to secrete a glycan with six or more residues in the repeating unit. Deleting any of the core genes causes a non-mucoid, EPS-phenotype, with the exception of *prmI* and *prmM*. *prmI* codes for a predicted Wzx-type repeating unit flippase that has no effect on colony morphology or EPS production. The apparent promiscuity of Wzx transporters can lead to genetic redundancy (39), and we predict that *prmI* is redundant with other glycolipid flippases in the LM7 genome. *prmM* codes for a predicted GT2-family glycosyltransferase. Deleting a GT that contributes to repeating unit assembly is expected to abolish polysaccharide production, but the Δ*prmM* mutant displays an unexpected phenotype. The colonies appear mucoid but do not take up Congo Red dye, suggesting that PrmM affects the properties of the EPS matrix but is not required for polysaccharide production.

Phenotypic analysis of *prm* genes that are not part of the predicted core pathway provide additional insight into the nuances of promonan biosynthesis. Deleting *prmB* (hypothetical protein) or *prmN* (GH10-family glycosylhydrolase) does not affect colony morphology or EPS production, indicating that these are dispensable for promonan biosynthesis under our growth conditions. *prmE* encodes a predicted AAA+-type ATPase that is required for promonan production. ATPases from the Bacterial Tyrosine Kinase (BYK) family participate in *wzy-* dependent polysaccharide production through a physical association Wzz family co-polymerases (such as PrmF) (43–45). However, *prmG* codes for a bonified BYK protein, and the identification of PrmE as a second ATPase that is required for polysaccharide production suggests that the architecture of the polymerization/export machinery may be more complex in LM7 than in previously described systems. *prmO* codes for an OafA-family acyltransferase that is required for promonan production. The implications of an acyltransferase being required for promonan production are discussed below.

Deleting *prmQ* causes a unique phenotype characterized by colonies that produce a Congo Red reactive matrix but show a rugose morphology distinct from the smooth, mucoid appearance of wild-type colonies (Fig 3B). *prmQ* codes for a predicted GH16-family glycosylhydrolase. These hydrolases are widespread across the tree of life. They display a range of carbohydrate modifying activities, with the majority of characterized representatives having polysaccharide endo-glycosidase activity (46). PrmQ has a predicted signal sequence for export to the periplasm, and we predict that PrmQ directly hydrolyzes pomonan during secretion, reducing the rigidity of the resulting polymers. The loss of such an activity would explain the rugose colony morphology seen in the Δ*prmQ* strain. A number of other polysaccharide biosynthesis clusters contain genes for glycosylhyrdolases (47, 48), and we suspect that their presence reflects a need to accommodate cell growth and division by modifying the extracellular matrix.

While we have not yet determined a detailed chemical structure of promonan, the identification of Glc, Gal and GlcA residues in their 3- and 4-linked forms indicates that the repeating unit contains six sugars in a linear, non-branching arrangement. A six-residue repeating unit might seem to align with the presence of six sugar transfer enzymes in the *prm* cluster. However, deleting the GT gene *prmM* does not alter colony mucoidy, but it causes cells to lose the ability to react with Congo Red dye. This phenotype suggests that *prmM* is not required for polysaccharide production but that its disruption alters the secreted matrix in some way. It is possible that PrmM incorporates a sugar into the growing repeating unit that is not required for downstream polymerization. Alternatively, deleting *prmM* could eliminate promonon production while activating the production of a separate, Congo Red insensitive extracellular matrix. Another intriguing aspect of promonan biosynthesis is the apparent requirement of an OafA-family acyltransferase gene (*prmO*) for polysaccharide production. The *oafA* gene from *Salmonella typhimurium* confers acetylation of an abequose residue in the O-antigen of the O5 serotype, but disruption of *oafA* simply abolishes O-antigen acetylation rather than eliminating polysaccharide production (49). The non-mucoid, EPS-phenotype of the Δ*prmO* mutant suggests a different role for this gene in the promonan pathway. Acylation of a glycolipid intermediate by PrmO must be required for assembly of a repeating unit that is competent for polymerization. In an analogous fashion, acetylation of a UPP-linked disaccharide by WcaF is required for assembly of the colonic acid repeating unit (50). Our composition analysis did not detect acylated sugars in the promonan extracts, but future studies aimed at determining chemical structures of intermediates in the pathway will help clarify the role of PrmO.

Analysis of functional redundancy among glycan assembly genes provided new insights into polysaccharide metabolism as a whole. Testing whether genes from different organisms could compensate for the loss of key enzymes in the *Caulobacter crescentus* holdfast pathway allowed us to dissect the substrate specificities of individual GTs. We identified a group of genes from diverse *Alphaproteobacteria* that appear to encode PHPT enzymes capable of generating UPP-Glc. We could also distinguish the specificities of WecG-family GTs from these organisms by showing that only a subset could compensate for the loss of *hfsJ*. While the precise reaction catalyzed by HfsJ has yet to be determined, our method efficiently delineated GTs that share its specificity. This cross-complementation approach provided molecular evidence that a pathway for producing holdfast-like adhesins is conserved in higher *Alphaproteobacteria*. More importantly, our results indicate that proteins with shared enzymatic activities but divergent sequences are interchangeable in polysaccharide biosynthesis pathways. Efforts to engineer the metabolism of other biopolymers have been hindered by the perturbation of protein-protein interactions and other unexpected disruptions when enzymatic functionalities are swapped genetically (51). *wzy*-dependent polysaccharide biosynthesis seems to present a truly modular system in which metabolic proteins can be interchanged without affecting pathway output.

Finally, this work highlights the extent to which bacterial polysaccharides remain under-sampled. Simply diluting water from a lake onto standard growth medium allowed us to identify a family of EPSs that is distinct from previously characterized polysaccharides. Promonan is synthesized through a relatively simple *wzy-*dependent pathway that utilizes housekeeping sugars. Nonetheless, we identified five GTs with unknown linkage specificities, an acyltransferase that likely acts on a glycolipid, a matrix modifying glycosylhydrolase and an ATPase that has not previously been associated with polysaccharide assembly. Detailed biochemical analysis of these proteins will continue to illuminate the molecular basis for generating the vast complexity among bacterial glycans. These novel enzymatic functions can also be used to engineer unnatural chemical complexity into bacterial polysaccharides through combinatorial biosynthesis. Characterizing novel polysaccharide pathways in detail not only adds to a growing collection of diversity generating enzymes, but streamlined assembly pathways like the one we identified in LM7 can also serve as scaffolds for genetic engineering of novel polymers.

## Materials and Methods

### Bacterial strains, growth conditions and genetic manipulations

Strains and plasmids used in this study are listed in Tables 5 and 6. Standard polymerase chain reaction (PCR), restriction digestion and Gibson assembly methods were used to develop plasmids. Strains, plasmids, primer sequences and details of construction are available upon request. *E. coli* was cultured in LB medium at 37°C supplemented with 1.5% (w/v) agar, 60mM diaminopimelic acid (DAP), 10 µg/mL tetracycline and 50 µg/mL kanamycin when necessary. *C. crescentus* was grown at 30°C in PYE medium supplemented with 1.5% (w/v) agar and 25 µg/mL kanamycin when necessary. Plasmids were introduced into *C. crescentus* by electroporation. *Sphingomonas sp.* LM7 was cultured at temperatures ranging from 18-30°C in either PYE medium or M2 medium with glucose. Agar (1.5% w/v), sucrose (3% w/v), tetracycline (5 µg/mL), kanamycin (25 µg/mL), cumate (50µM), Congo Red (20µg/mL) and Coomassie Blue (10µg/mL) were added to LM7 cultures when necessary. Plasmids were introduced into LM7 by conjugation with WM3064 as the donor strain. Gene deletions were generated using a two-step approach with *sacB-*based counterselection.

**Table 5:** Plasmids used in this study.

| Plasmid | Description | Antibiotic | Reference |
| --- | --- | --- | --- |
| pKMW3 | Plasmid for delivery of barcoded himar transposon | Km | (53) |
| pNPTS138 | Suicide plasmid for making unmarked deletions in *Sphingomonas sp. LM7*; carries *sacB* for counter-selection | Km | M. R. Alley |
| pDH298 | To delete *prmJ*; Gibson cloning of fused upstream and downstream regions of BXU08_RS00395 | Km | This work |
| pDH354 | To delete *wzyB*; Gibson cloning of fused upstream and downstream regions of BXU08_RS19300 | Km | This work |
| pDH1289 | To delete *prmA*; Gibson cloning of fused upstream and downstream regions of BXU08_RS00350 | Km | This work |
| pDH1290 | To delete *prmB*; Gibson cloning of fused upstream and downstream regions of BXU08_RS00355 | Km | This work |
| pDH1291 | To delete *prmC*; Gibson cloning of fused upstream and downstream regions of BXU08_RS00360 | Km | This work |
| pDH1292 | To delete *prmD*; Gibson cloning of fused upstream and downstream regions of BXU08_RS00365 | Km | This work |
| pDH1293 | To delete *prmE*; Gibson cloning of fused upstream and downstream regions of BXU08_RS00370 | Km | This work |
| pDH1294 | To delete *prmF*; Gibson cloning of fused upstream and downstream regions of BXU08_RS00375 | Km | This work |
| pDH1295 | To delete *prmG*; Gibson cloning of fused upstream and downstream regions of BXU08_RS00380 | Km | This work |
| pDH297 | To delete *prmH*; Gibson cloning of fused upstream and downstream regions of BXU08_RS00385 | Km | This work |
| pDH1296 | To delete *prmI*; Gibson cloning of fused upstream and downstream regions of BXU08_RS00390 | Km | This work |
| pDH1297 | To delete *prmK*; Gibson cloning of fused upstream and downstream regions of BXU08_RS00400 | Km | This work |
| pDH1298 | To delete *prmL*; Gibson cloning of fused upstream and downstream regions of BXU08_RS00405 | Km | This work |
| pDH1299 | To delete *prmM*; Gibson cloning of fused upstream and downstream regions of BXU08_RS00410 | Km | This work |
| pDH1300 | To delete *prmN*; Gibson cloning of fused upstream and downstream regions of BXU08_RS00415 | Km | This work |
| pDH1301 | To delete *prmO*; Gibson cloning of fused upstream and downstream regions of BXU08_RS00420 | Km | This work |
| pDH1302 | To delete *prmP*; Gibson cloning of fused upstream and downstream regions of BXU08_RS00425 | Km | This work |
| pDH331 | To delete *prmQ*; Gibson cloning of fused upstream and downstream regions of BXU08_RS00430 | Km | This work |
| pDH1303 | To delete *rfbA*; Gibson cloning of fused upstream and downstream regions of BXU08_RS19140 | Km | This work |
| pDH1248 | pQF: pCM62 with cymR*, PQ5, and MCS for N- and C-terminal fusions to 3FLAG tag; Tcr | Tc | (26) |
| pDH1304 | pQN: pQF derivative with N-terminal FLAG tagging region removed; used for complementation experiments | Tc | This work |
| pDH1305 | Gibson cloning of fused upstream and downstream regions of BXU08_RS00385 into cumate inducible plasmid pQN | Tc | This work |
| pDH1306 | Gibson cloning of fused upstream and downstream regions of BXU08_RS00430 into cumate inducible plasmid pQN | Tc | This work |
| pDH1307 | Gibson cloning of fused upstream and downstream regions of BXU08_RS00395 into cumate inducible plasmid pQN | Tc | This work |
| pDH118 | pMT585; pMTLS4259: Integrating vector for xylose induceable expression of C-terminally GFP tagged proteins in *Caulobacter* | Km | (57) |
| pDH444 | Gibson cloning of *C. crescentus* CB15 CC_2425 into pDH118 | Km | This work |
| pDH478 | Gibson cloning of *R. leguminosarum* RL1661 into pDH118 | Km | This work |
| pDH477 | Gibson cloning of *B. ovis* BOV_0775 into pDH118 | Km | This work |
| pDH445 | Gibson cloning of *P. inhibens* PGA1_262p00380 into pDH118 | Km | This work |
| pDH1288 | Gibson cloning of *Shingomonas* sp. LM7 BXU08_RS00350 into pDH118 | Km | This work |
| pDH476 | Gibson cloning of *E. coli* wcaJ into pDH118 | Km | This work |
| pDH124 | Gibson cloning of *C. crescentus* CB15 CC_0095 into pDH118 | Km | This work |
| pDH125 | Gibson cloning of *B. ovis* BOV_A0095 into pDH118 | Km | This work |
| pDH128 | Gibson cloning of *R. leguminosarum* RL0741 into pDH118 | Km | This work |
| pDH295 | Gibson cloning of *Sphingomonas* sp. LM7 BXU08_RS00385 into pDH118 | Km | This work |
| pDH155 | Gibson cloning of *E. coli* wecG into pDH118 | Km | This work |

**Table 6:**
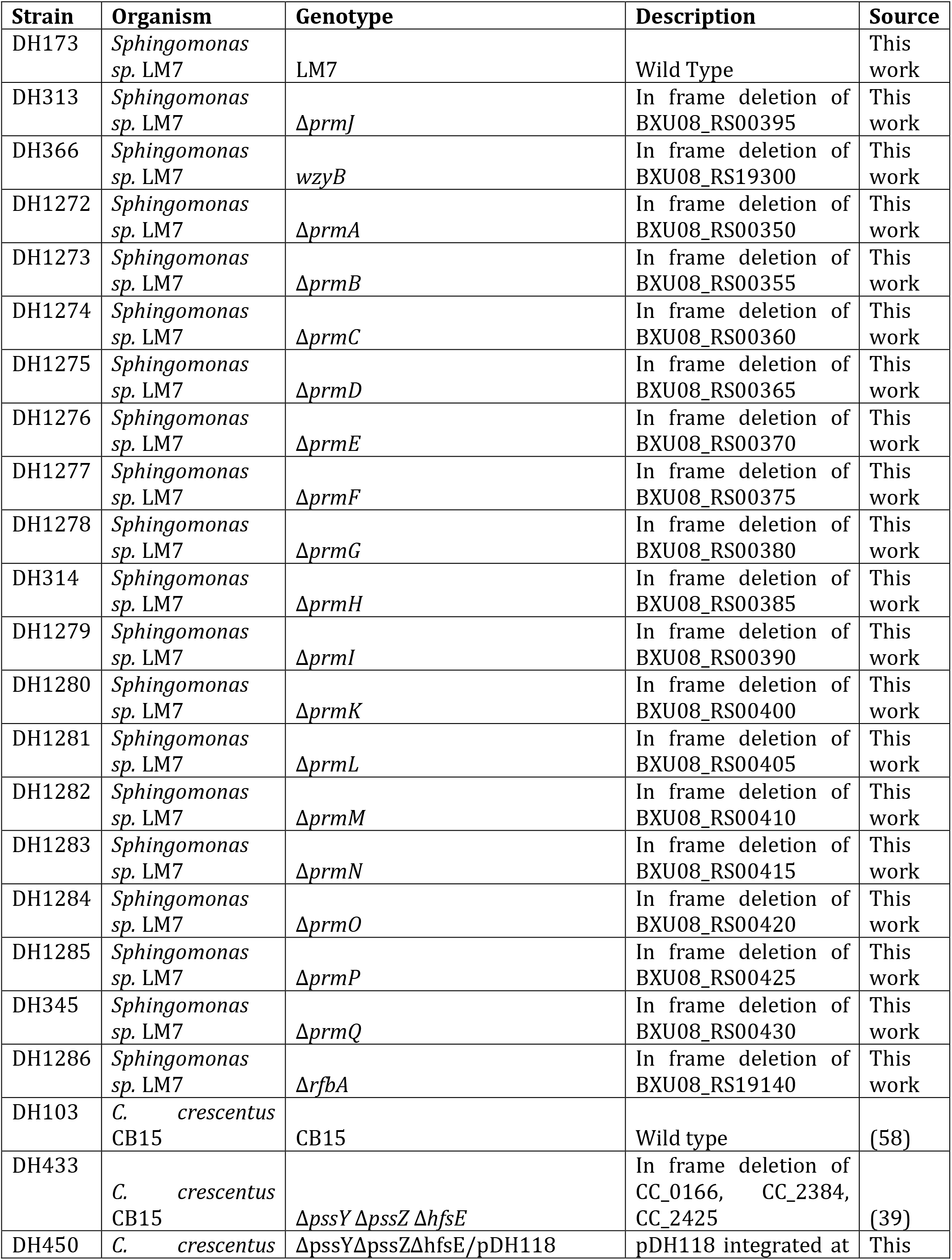
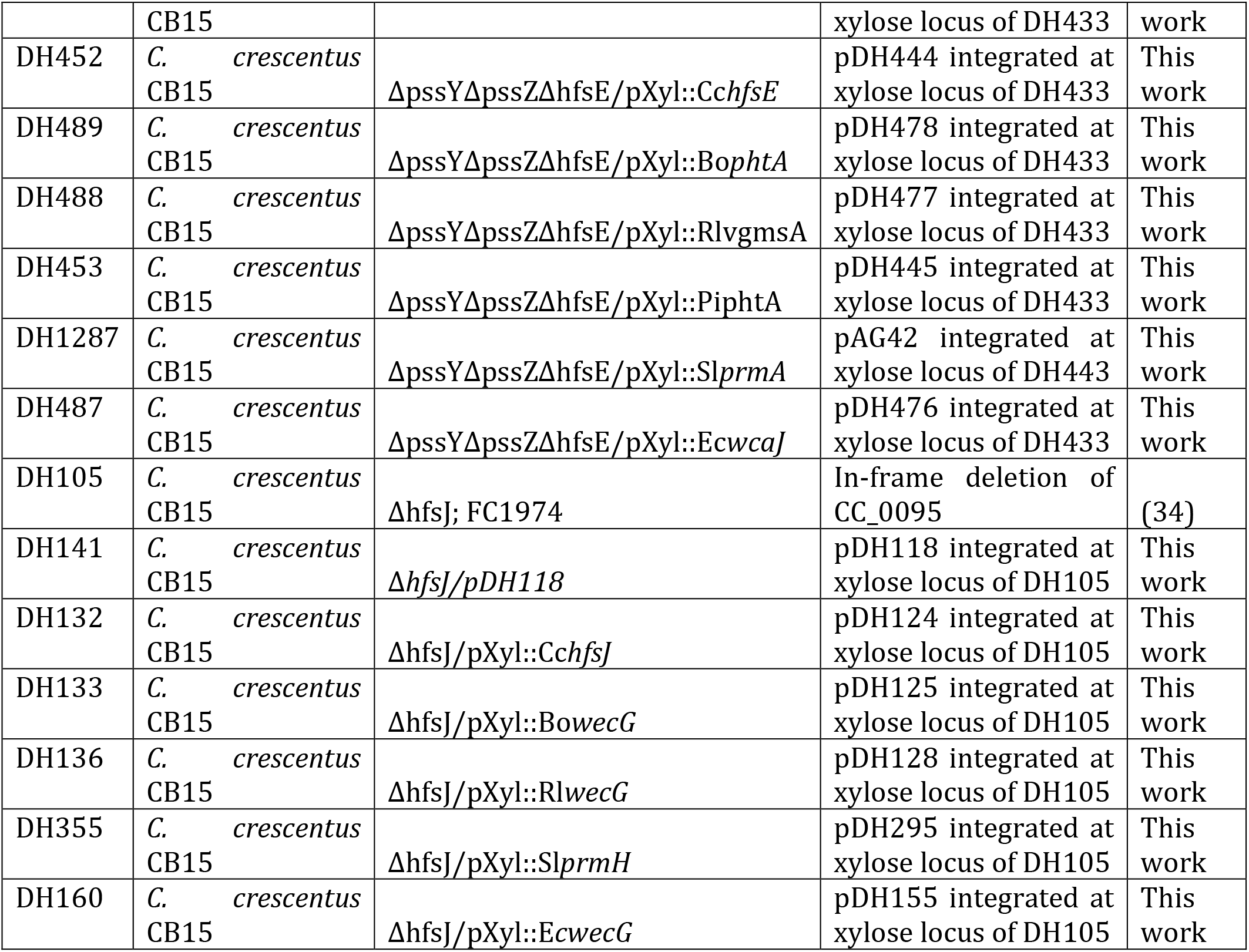
Strains used in this study.

### Isolation of Sphingomonas sp. LM7

Water from Lake Michigan was collected near the shore of Promontory Point in Chicago, IL using a plastic beer pitcher attached to the end of a 10ft metal pole. The water was serially diluted, plated on PYE agar and incubated at 30°C. A single, yellow colony was isolated for further study.

### Genome sequencing

Genomic DNA isolated from LM7 was sequenced on a PacBio RS II instrument at the University of Wisconsin – Milwaukee Great Lakes Genomics Center. Genome assembly using HGAP3 (52) yielded a single, circular contig based on ∼228x coverage. The genome was deposited under NCBI RefSeq NZ_CP019511.1.

### Transposon mutagenesis and analysis of mutants with altered colony morphology

*Sphingomonas sp.* LM7 was mutagenized with a modified mariner transposon. WM3064 cells carrying pKMW3 (53) were mixed with LM7 cells at a ratio of approximately 1:4. The mixture was spotted on PYE plates containing DAP and incubated overnight at room temperature. Cells from the conjugation reaction were resuspended in PYE, diluted appropriately and spread onto 150mm plates containing M2 medium with 2% (w/v) glucose and kanamycin. Plates were incubated for 5 days at room temperature. Mutants were screened visually for changes in colony morphology. Mutants of interest were isolated and patched onto PYE agar containing kanamycin for further study.

Mutations were mapped using a two-step, arbitrary PCR strategy. 2x GoGreen Master Mix (Promega) was used for these reactions. A primer containing a sequence from the 3’ end of the transposon cassette (U1 fw: 5’-GATGTCCACGAGGTCTCT-3’) and a primer containing the M13 priming sequence fused to a random heptamer (M13-N_7_: 5’ TGTAAAACGACGGCCAGTNNNNNNN-3’) were used to amplify the junction between the transposon and genomic insertion site. The conditions for this reaction were as follows: 95°C for 2min, 35 cycles of 95°C for 30sec, 38°C for 30sec and 72°C for 1min, followed by a 5min hold at 72°C. The reaction was treated with ExoSap-IT (ThermoFisher), and a second, nested amplification reaction was then performed using this ExoSap treated product as a template. A different region from the 3’ end of the transposon cassette (U2out: 5’ CGTACGCTGCAGGTCGAC-3’) and the M13 priming sequence served as primers for this second reaction. Reactions were incubated at 95°C for 2min, and a 40 cycle touchdown program of 95°C for 30sec, a 30s annealing step and 72°C for 1m was applied followed by a 5min incubation at 72°C. The annealing step was performed at 68°C for the first cycle and decreased 0.5°C with each cycle. The nested reaction products were treated with ExoSap and subjected to Sanger sequencing to identify the genomic sites of transposon insertion.

### Isolation of Promonan EPS

Overnight cultures of *Sphingomonas sp.* LM7 grown in PYE, were used to inoculate M2 medium supplemented with 2% (w/v) D-glucose. Cultures were growth at 18°C for 72 hours with shaking at 200rpm. Cells were removed from the culture broth by centrifugation at 8000 x g, and the resulting supernatant was transferred to fresh tubes. Two volumes of ethanol was added to the spent medium, and the mixture was incubated overnight at 4°C. Precipitate was harvested by centrifuging for 1hr at 8000 x g. After discarding the supernatant, the gel-like insoluble fraction was resuspended in water. Tris-HCl pH 7.4, MgCl_2_ and turbonuclease were added to final concentrations of 10mM, 2mM and 1µg/mL, respectively. The nuclease digest was incubated overnight at 37°C. Tris-HCl pH 8.5 and proteinase K were then added to final concentrations of 25mM and 100µg/mL, respectfully, and the digest was again allowed to proceed overnight at 37°C.

An equal volume of aqueous phenol was added to the extraction. The mixture was homogenized thoroughly, 10-15 drops of chloroform was added and the layers were allowed to separate for 30min. The upper aqueous layer was then transferred to dialysis tubing (8kDa cutoff) and was dialyzed against deionized water for 72 hours. The dialysate was transferred to fresh deionized water every 24hrs. The resulting extracts were transferred to glass tubes, incubated at - 20°C overnight and lyophilized to dryness.

### Monosaccharide composition analysis

The analysis was performed by combined gas chromatography-mass spectrometry (GC-MS) of the O-trimethylsilyl (TMS) methyl glycoside derivatives produced from the sample by acidic methanolysis. First, a 1mg/mL solution of each sample was made. Of this solution, 200μL was combined with 2μL of a 10mg/mL stock solution of myo-inositol, and the resulting mixture was lyophilized. To the dried samples was added 15 drops of 1M methanolic-HCl. The samples were incubated for 18h at 80°C. The following day, the samples were removed from the heating block and cooled to room temp. Once cooled, they were dried with dry nitrogen gas, during which 5 drops of MeOH was added every 3 min. Next, the samples were N-acetylated with 8 drops of MeOH, 4 drops of pyridine, and 4 drops of acetic anhydride, and the mixture was incubated at 100°C for 30 min. Once cooled to room temperature, the samples were dried with dry nitrogen gas. Five drops of MeOH were added after 5 min, and after complete drying, 10 drops of Tri-Sil HTP reagent were added to each sample, followed by incubation at 80°C for 20 min. After evaporation of the solvent, 100μL of hexane was added to each sample. The samples were briefly vortexed and centrifuged, and then transferred to GC vials. For analysis, 1μL was injected. GC-MS analysis of the TMS methyl glycosides was performed on an AT 7890A GC interfaced to a 5975B MSD, using an EC-1 fused silica capillary column (30 m x 0.25 mm ID). The temperature gradient involved an initial 2min hold at 80°C, a 20°C/min ramp to 140°C, a 2min hold at 140°C, a 2°C/min ramp to 200°C, a 30°C/min ramp to 250°C, and a final 5min hold at 250°C.

### Glycosyl linkage analysis

The identification of glucuronic acid in the promonan extracts complicated the analysis of partially-methylated alditol acetates (PMAAs) because uronic acids diminish solubility in dimethylsulfoxide, leading to poor results. We deployed a recently developed method (25) whereby the acidic polysaccharide is first acetylated in ionic liquid to increase solubility, leading to vastly improved recoveries.

Samples were acetylated by dissolving in 300μL of the ionic liquid 1-ethyl-3-methylimidazodium acetate and sonicating to fully dissolve. Once fully dissolved, 500μL of acetic anhydride and 50μL of 1-methylimidazole were added. The samples were then stirred for 10min. After stirring, 2mL of dichloromethane (DCM) and 2mL of H_2_O were added. Each sample was vortexed and centrifuged briefly, and then the aqueous layer was carefully discarded. Each sample contained a white interphase. All of the water was not removed so as not disturb this phase. The extraction was repeated 4 times. On the final wash, the organic (DCM) layer was removed and dried down with dry nitrogen. Once the DCM was completely removed, the samples were lyophilized. After this, the samples were dissolved in 300μL of dimethylsulfoxide (DMSO). Next, 300μL of potassium methylsulfinylmethylide in DMSO (∼1M) was added to each sample, which was stirred for 1hr. The samples were then placed in an ice bath until the solution solidified. 100μL of iodomethane was then added at such a rate to keep the solution from boiling. Once the solution color changed from dark brown to light brown, another 100µL of iodomethane was added, and the samples were stirred for about 25min to ensure full permethylation of the uronic acids. Another DCM extraction was performed, and the DCM layer was dried with dry nitrogen after the final wash. The samples were then lyophilized overnight before continuing with reduction.

To begin reduction, 300μL of a 10mg/mL solution of lithium aluminum deuteride (LiAlD_4_) in tetrahydrofuran (THF) was added to the dried sample, which was then incubated at 80 °C for 4 h. The reaction was then neutralized with 2 drops of acetic acid (AcOH). Five drops of MeOH were added to each. The samples were then dried with dry nitrogen gas. Once fully dry, 10 drops of 9:1 MeOH:AcOH was added and the samples were dried again. The 10 drops of 9:1 MeOH:AcOH was added and dried one more time. Next, 10 drops of MeOH were added and dried with dry nitrogen, and this step was repeated two times. The samples were redissolved in 2 mL of H_2_O, transferred to 6-kDa dialysis bags, and dialyzed against 2 exchanges of DI water per day for 2 days. Following dialysis, the samples were permethylated according to the following procedure.

Sodium hydroxide (NaOH) base was prepared according to the procedure of Anumula and Taylor (54), and of this, 300µL was added to all samples. The sample containing base was stirred for 15 minutes. A volume of 150µL of iodomethane was added, and the mixture was stirred for 25min. The steps adding base and iodomethane were repeated one more time to ensure that the samples were fully methylated. Once mixing was completed, 2 mL of dichloromethane (DCM) and 2 mL H_2_O were added. The tube was vortexed and centrifuged briefly, then the upper (aqueous) layer was removed. This step was repeated four more times. After the final wash, the DCM layer was removed into a separate tube and dried with dry nitrogen gas.

The samples were hydrolyzed in 2M trifluoroacetic acid (TFA) for 2 h in a sealed tube at 120 °C. The hydrolyzed samples were cooled to room temperature, and the TFA was evaporated using dry nitrogen gas. Once fully dry, 10 drops of isopropanol (IPA) were added. The samples were dried again, and the addition and drying of IPA was repeated two times. On the last addition of IPA, a 10-mg/mL solution of NaBD_4_ and 1 M NH_4_OH was made with 9.48 mg of NaBD_4_ and 948µL of NH_4_OH. Of the NaBD_4_ solution, 400µL was added to each sample. The samples were left to incubated for reduction overnight. The following day, the basic reaction was neutralized with 3 drops of acetic acid (AcOH). Five drops of MeOH were added to each. The samples were then dried with dry nitrogen gas. Once fully dry, 10 drops of 9:1 MeOH:AcOH was added and the samples were dried again. Then, 10 drops of 9:1 MeOH:AcOH was added and dried one more time. Next, 10 drops of MeOH were added and dried with dry nitrogen, and this step was repeated two times. The samples were then O-acetylated with 250µL of acetic anhydride and 230µL of TFA, incubated at 50 °C for 10min. Once cooled to room temperature, a DCM extraction was done by adding 2 mL of DCM and 2 mL of H_2_O to each sample. The mixture was briefly vortexed and centrifuged. The aqueous layer was carefully decanted, and the process was repeated 4 times. On the final wash, the DCM layer was removed into a separate tube and dried with dry nitrogen. Five drops of DCM were then added and transferred to GC vials. 1µL of each was injected into the GC-MS for analysis. The resulting linkages were analyzed on an Agilent 7890A GC interfaced to a 5975C MSD, electron impact ionization mode. Separation of natural monosaccharides was performed on a 30-m Supelco SP-2331 bonded phase fused silica capillary column. The column was subjected to a 1min initial hold at 60°C. The temperature was then increased to 170°C over 4min, raised to 235°C over 16.5min, held at 235°C for 2min and finally raised to 240°C over 1.4min followed by a 12min final hold at 240°C.

### Crystal violet staining assay

Crystal violet staining analysis of adhesion in *C. crescentus* strains was performed as described previously (55). Overnight cultures grown in PYE were normalized to an OD_600_ of 0.5, and 1μL of normalized culture was added to 450μL of PYE in a 48-well microtiter plate. Plates were incubated for 18-20hrs at 30°C with shaking at 155rpm. Culture broth was discarded, the wells were washed thoroughly with tap water and 500μL of 0.01% (w/v) crystal violet was added to each well. The plate was shaken at 155rpm with the dye for 5min before the dye was discarded and the wells again washed thoroughly with tap water. Retained dye was then dissolved in 500uL of ethanol by shaking for 5min at 155rpm. Absorbance at 575nm was measured in each well.

### Visualization of holdfast production

Overnight cultures of *C. crescentus* strains of interest were grown overnight, diluted into fresh PYE and allowed to reach mid-log phase (OD_600_=0.4-0.5). 200uL of culture was added to a clean tube, 1uL of Alexa647-conjugated wheat germ agglutinin (fWGA) was added from a 2mg/mL stock and the labeling reaction was incubated at room temperature for 5min in the dark. 1mL of sterile water was added to the tube and the mixture was centrifuged for 2min at 6000 x g to sediment cells. The supernatant was discarded, cells were resuspended in the residual liquid and 1uL was spotted onto 1% agarose pads. Microscopy was performed using a Nikon Ti-E inverted microscope equipped with an Orca Fusion BT digital CMOS camera (Hamamatsu). Fluorescence images were collected using a Prior Lumen 200 metal halide light source and an mCherry-specific filter set (Chroma).

## Data availability

Reads and metadata from the LM7 genome sequencing project can be accessed through NCBI BioProject PRJNA363095. Sanger sequencing reads from transposon insertion mapping, microscopy images and any other data collected during this work have been archived and are available upon request.

## Acknowledgements

We would like to thank Sean Crosson, Aretha Fiebig and Swaine Chen for assistance with the LM7 genome assembly. This work was supported by a Beckman Young Investigator award to D.M.H., National Institutes of Health award R35GM150652 to D.M.H. and startup funds from the University of Wisconsin – Madison to D.M.H. This work was supported by the U.S. Department of Energy, Office of Science, Basic Energy Sciences, Chemical Sciences, Geosciences and Biosciences Division, under award #DE-SC0015662 to P.A.

